# Dramatic shape changes occur as Cytochrome c folds

**DOI:** 10.1101/2020.06.25.171926

**Authors:** Serdal Kirmizialtin, Felicia Pitici, Alfredo E Cardenas, Ron Elber, D. Thirumalai

## Abstract

Extensive experimental studies on the folding of Cytochrome c (Cyt c) make this small protein an ideal target for atomic detailed simulations for the purposes of quantitatively characterizing the structural transitions and the associated time scales for folding to the native state from an ensemble of unfolded states. We use previously generated atomically detailed folding trajectories by the Stochastic Difference Equation in Length (SDEL) to calculate the time-dependent changes in the Small Angle X-ray scattering (SAXS) profiles. Excellent agreement is obtained between experiments and simulations for the time dependent SAXS spectra, allowing us to identify the structures of the folding intermediates, which shows that Cyt c reaches the native state by a sequential folding mechanism. Using the ensembles of structures along the folding pathways we show that compaction and the sphericity of Cyt c change dramatically from the prolate ellipsoid shape in the unfolded state to the spherical native state. Our data, which provides unprecedented quantitative agreement with all aspects of time-resolved SAXS experiments, shows that hydrophobic collapse and amide group protection coincide on the 100 microseconds time scale, which is in accord with ultrafast Hydrogen/Deuterium exchange studies. Based on these results we propose that compaction of polypeptide chains, accompanied by dramatic shape changes, is a universal characteristic of globular proteins, regardless of the underlying folding mechanism.

## Introduction

Major advances in experiments have elucidated the complexities of the folding process,^1–6^ thus providing an impetus to develop computational methods for making quantitative predictions. In addition, theory and simulations using a variety of coarse-grained models have provided a framework for anticipating different folding scenarios.^7–11^ Despite the conceptual advances, it still remains a challenge to predict the folding pathways of specific proteins under conditions that are typically used in experiments.^12^ There are compelling reasons to rise to this grand challenge. First, advances in single molecule experiments are starting to provide detailed information about the changes in the ensemble of conformations during the folding reaction as the external conditions (temperature, mechanical force, or denaturant concentrations) are varied.^1,13,14^ Secondly, fast mixing methods in conjunction with small angle X-ray (SAXS) measurements yield global information about the changes in the size of a protein and the extent to which secondary structure is formed during the collapse process.^15,16^ The results from these experiments and related methods^17–19^ demand precise computations for specific systems so that one can predict the molecular details of the folding process. Although recent advances in computer hardware and other technological developments have been used to generate long folding trajectories using all atom molecular dynamics simulations,^20–23^ to date they have not yielded results that can be quantitatively compared with experiments, except in rare cases. Here, we show that the time-resolved SAXS profiles calculated using folding trajectories for Cytochrome c (Cyt c) are in quantitative agreement with experiments. With this validation, we establish that the changes in the shapes of Cyt c, which folds in a sequentially manner, are dramatically going from a prolate ellipsoid to a sphere. Without adjusting any parameter, our simulations produce near quantitative agreement with hydrogen/deuterium exchange experiments, in the process revealing the molecular details in the early stages of Cyt c folding. By monitoring the Trp59-Heme distance in the early stages of folding from simulations, we explain the origins of the apparent differences in the interpretation regarding the kinetic barrier to Cyt c collapse.

Time resolved SAXS experiments have provided details of the relationship between the changes in the radius of gyration (*R*_*g*_) and the secondary structure formation setting the stage for two issues that we address here. The first is concerned with changes in the shape of the protein as it folds, the importance of which can be appreciated based on the following arguments. The *R*_*g*_ values in the denatured states scales as *R*_*g*_ *≈ a*_*D*_*N*^*ν* 24^ where the Flory exponent *ν ≈* (0.5 *−* 0.6), *N* is the number of amino acid residues, and *a*_*D*_ is a characteristic length. Folded states of proteins are nearly maximally compact with 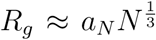 where *a*_*N*_ *≈* 3Å.^25^ Thus, during the folding process, the polypeptide chain must become compact.^26–30^ The shape of the unfolded state is expected to be a prolate ellipsoid while the native state is spherical.^25^ It follows that the transition from the unfolded to the native structure involves not only a large surface area change but also a dramatic change in the overall shape (prolate ellipsoid to a sphere). The second issue concerns the early stages of folding. In some proteins, compaction and acquisition of native structure occur concurrently,^26^ a prediction that has been validated using experiments.^31^ Theoretical considerations and simulations of simple protein models show, in accord with time resolved SAXS experiments on several proteins, that the collapse process occurs in three distinct stages with most of the changes in *R*_*g*_ occurring relatively early in the folding reaction.^26^ For example, the 104 amino acid residue Cytc (the protein studied here) collapses rapidly to a compact intermediate globular state^15,16^ in perhaps as little as 50*µs*. The rapid collapse is followed by a slower phase that leads to the formation of secondary structures, which raises the question of what is the relationship between chain compaction, secondary structure formation, and folding in Cyt c?

## Methods

### Simulations

We used the publicly distributed molecular dynamics package MOIL,^32^ which uses a combination of AMBER and OPLS force fields. The Stochastic Difference Equation in Length (SDEL) requires a suitable start and end points, which in the case of protein folding correspond to unfolded and folded states, respectively. The folded structure was generated by minimizing the energy of the native structure (taken from the Protein Data Bank with code 1HRC) after removing the crystallographic water molecules. We used the Generalized Born model^33^ to account for the solvation energy. The model includes covalent binding of the heme group to Cys14 and Cys17, but does not have coordination of the heme iron to Met90. The unfolded structures were initially generated at a high temperature, and after a sequence of energy minimization an ensemble of starting structures were obtained at the simulation temperature of 300 *K*. Additional details can be found below and in the methods section of a previous study.^34^

### Computing the SDEL Trajectories

The computations are based on the folding pathways generated by Cardenas and Elber^34^ using the SDEL methodology. The simulation method used to generate the folding trajectories for the present manuscript was also used in the successful study of the folding of protein A,^35^ which provides additional justification for the simulation method. Each trajectory, parameterized by length *l*, that is obtained by solving a boundary value problem. Given the two end structures (the folded conformation, *X*_*f*_, and another *X*_*u*_, that is sampled from the set of unfolded conformations generated at finite temperature as described below), we optimize the functional (action) 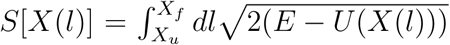 for the entire trajectory. Here, *X*(*l*) is the coordinate vector of an all atom model of Cyt c, parameterized as a function of the arc-length *l, E* is the total energy, and *U*(*X*(*l*)) is the potential energy including the part that accounts for solvent effects. In our calculations, a trajectory is represented by a discrete set of coordinates *X*_*i*_ with *i* = 1, 2, …*N* that are equally separated along the path by |*X*_*i*_ *− X*_*i*+1_| with *i* = 1, 2, ..(*N −* 1). Using an implicit solvation model,^33^ we generated 26 folding trajectories that start from an ensemble of unfolded structures and end at the native conformation. From each SDEL trajectory we harvested 900 structures, which were used to compute the time-dependent observables.

### Generation of the Unfolded State Ensemble

To create an ensemble of unfolded structures, we used high temperature MD simulation. To ascertain if residual structure persists in the unfolded state we performed molecular dynamics simulations with a modified non-bonded interactions that include only a repulsive term (no electrostatic forces or attractive van der Waals energies). Such a model mimics a self-avoiding polymer that retains the covalent geometry of Cyt c. These simulations were done at the temperature of 1,500 *K* for 2 nanoseconds. We picked 200 structures with the root mean square distance of at least 6 Å away from each other and minimized conformations with the full non-bonded interactions using Generalized Born term for solvation. The minimization fixes local geometries and remove bad contacts but it also changes a little the overall shape of the protein, which is our prime concern here. The structures were used to calculate the scattering function for Cyt c in the limit when the interactions are determined only by excluded volume.

### Calculation of the Time-resolved Small Angle X-Ray Scattering (SAXS) Spectra

The 900 structures from each of the 26 trajectories connecting the folded and unfolded states used to calculate the scattering function *I*_*α*_(*q*|*l*_*j*_) as a function of the momentum transfer, 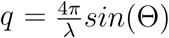 *sin*(Θ) (*α* is the trajectory label, *l*_*j*_ is the arc length of the SDEL path at up to configuration *j, λ* is the wavelength and 2Θ is the scattering angle). The scattering functions were calculated using the CRYSOL program^36^ with explicit averaging over all possible orientations to ensure that spectra are isotropic, making it appropriate for comparison with the solution phase SAXS profiles. We averaged the scattering function over all different trajectories at a given path length *l*_*j*_ using 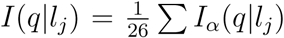. We implicitly assume that the path length is monotonic with trajectory times, an assumption which we later examined for consistency.

Using the crystal structure of the Cyt c we first fit the computationally generated SAXS profile for the native state ensemble with the experimental SAXS curve representing the folded state. For the purposes of comparing to the SAXS experiments,^16^ we normalized the Kratky plot by dividing all the spectra by the maximum in *q*^2^*I*(*q*) (*I*(*q*) is the scattering intensity at the wave vector *q*) corresponding to the experimentally measured value in the native state. This allowed us to find the normalization constant between experiments and simulations. Note that the normalization could also be calculated based on *I*(*q* = 0). However, the experimental data was not available for zero angle scattering. In addition, extracting *I*(*q* = 0) from Kratky plots creates uncertainties. Therefore, we used the Kratky plots for direct comparison. We used the same constant for all other time intervals assuming that the protein concentration stayed constant during measurements.

In the SAXS experiments,^16^ the authors presented only 10 out of the total 33 SAXS spectra at different folding times. We digitized the ten spectra and compared them with the selected spectra from Kratky plots obtained from simulations.

To match the simulated *I*(*q*|*l*_*j*_) and measured *I*(*q*|*t*)^16^ requires a mapping between *l*_*j*_ and real time *t*. For this purpose averaged scattering function at a given path length is compared with the real time *I*(*q*|*t*). The path length *l*_*j*_ that gives the best agreement to the experimental curve is selected. The remarkable linear relationship between the arc length and the logarithm of physical time is obtained (Figure 1 inset), allowing us to convert *I*(*q*|*l*_*j*_) to *I*(*q*|*t*). This agreement makes it convenient to match the simulation and experimental results, without any additional adjustments.

**Figure 1:**
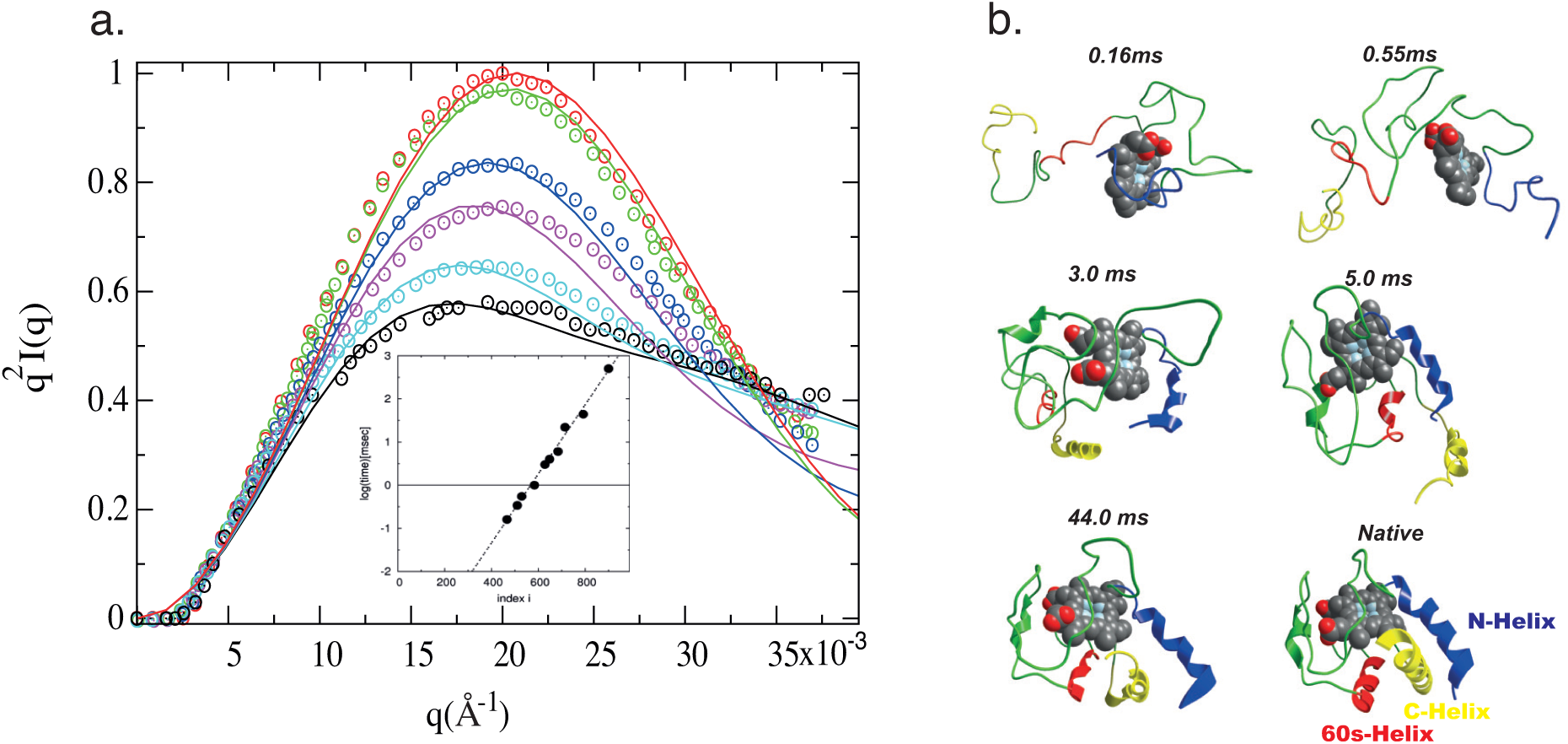
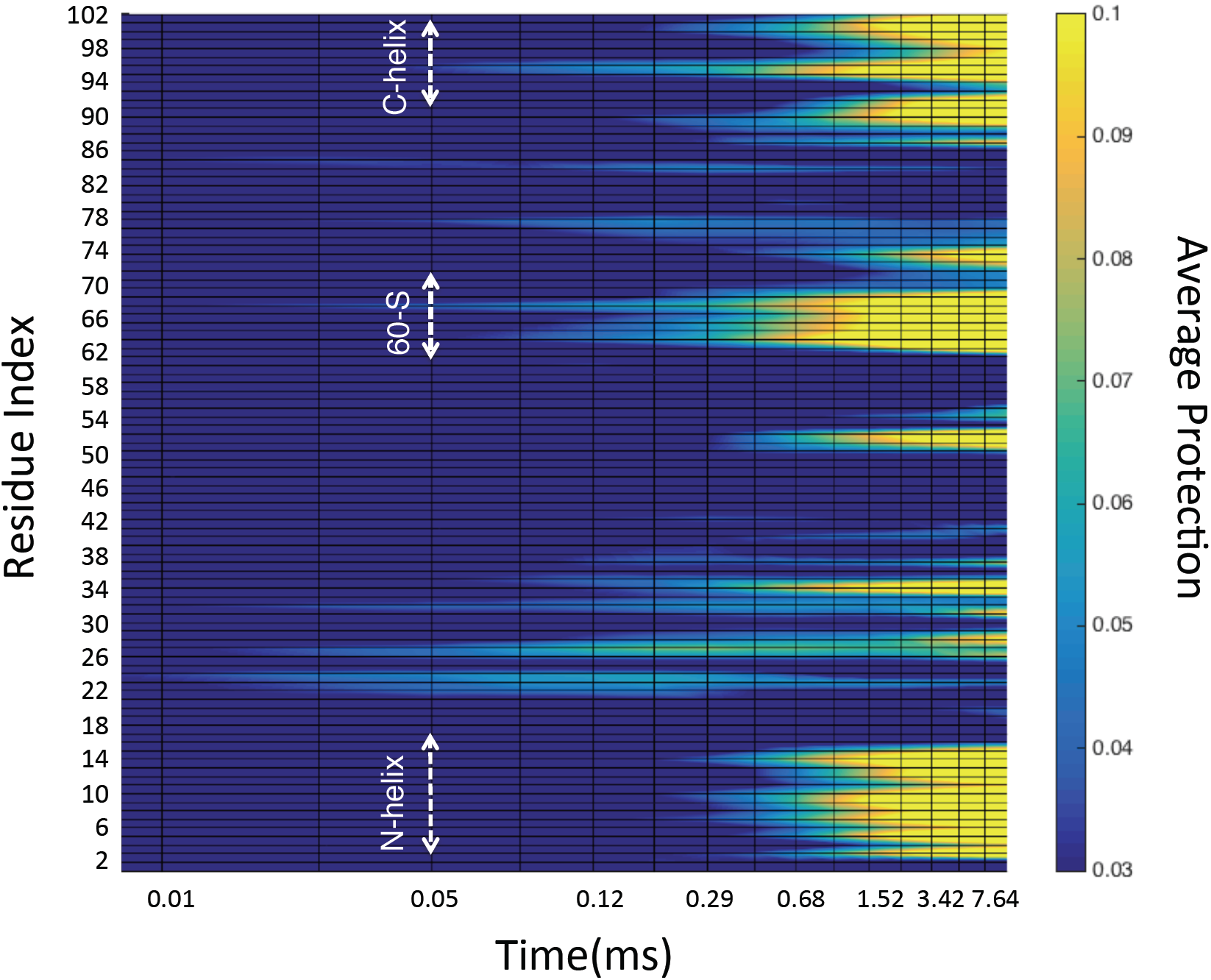
(a) Time resolved Kratky plots during the folding of Cyt c. Open circles are taken from the SAXS measurements,^16^ and solid lines are simulation results. Colors represent different times during the folding process; black (*t* = 160*µs*), cyan (*t* = 550*µs*), purple (*t* = 3*ms*), blue (*t* = 6*ms*), green (*t* = 44*ms*), and red is the native state. The inset shows the relationship between experimentally measured time and SDEL path index i, which is given by *t* = 0.08*i* − 4.5361. (b) Representative structures during folding. Conformations are chosen from among the 26 paths that best represent the Kratky plots in Fig. 1a. Images are created with MOIL.^32^ (c) Protection of each amide group of Cyt c as a function of time (see SI for details) for comparison to the ultra-fast time resolved H/D exchange experiments.^43^

### Calculation of the Amide Group Protection

It would be most interesting to predict the propensity of amide hydrogen for exchange with deuterium from simulations. It is found that amide group protection is empirically correlated with the presence of hydrogen bond of the backbone amide nitrogen to an acceptor.^37–39^ The protection factors are most directly correlated with solvent accessible surface area (SASA) of the amide group.^40–42^ We found that SASA based amide group protection describes the folded state of cyt c when compared with experiments.^43^ Because we used implicit solvent in the simulations it is most appropriate to use SASA as a proxy for the protection factor.

For that purpose, to study the amide group protection during folding we compute the surface area change as a function of time.

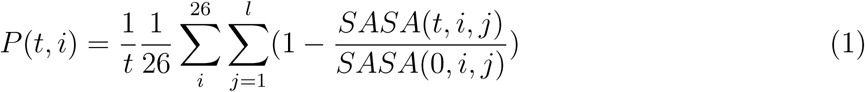

where, *SASA*(*t, i, j*) is the solvent accessible surface area (SASA) of the amide group of residue *j* of trajectory *i* at time *t. SASA*(0, *i, j*) is its value at the unfolded state (*t* = 0). The ratio in the summation changes in the range (0,1) where 0 corresponds to no change in the protection of the amide group with respect to the unfolded state. The lower the *SASA*(*t, i, j*) the more the protection of the amide hydrogens, hence slower the exchange rates. We calculated SASA using the algorithm^44^ implemented in Gromacs.^45^

### Order Parameters Characterizing the Shapes

To quantify the changes in the shape of Cyt c we used three global structural measures, which were constructed using the eigenvalues of the tensor of inertia *λ*_*i*_ (*i* = 1, 2, 3). The radius of gyration, 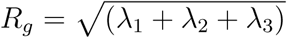 gives the size of the protein. The extent of deviation from spherical symmetry can be expressed in terms of the asymmetry parameter,

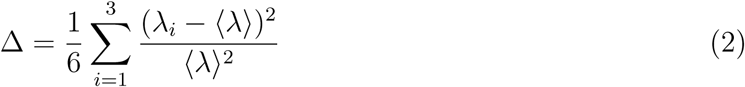

Where 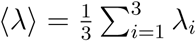. The shape parameter, S,

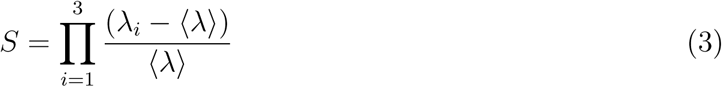

allows us to characterize the global shape of the polypeptide chain as either prolate (*S >* 0) or oblate (*S <* 0) ellipsoid. The values Δ lies between zero and unity and *S* ranges from −0.25 to 2.0. For a sphere, both parameters are zero, while for a rod, Δ = 0.25 and *S* = 0.2.

### Calculations of Fluorescence Energy Transfer

To compare the simulated folding pathways with experiments, we calculated the fluorescence energy transfer (FRET) from the distance, *R*, between Trp59 to the covalently attached heme group using,

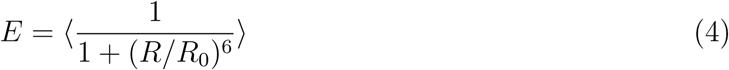

where the averag is over the folding trajectories, *R*_0_ is the Förster radius for the donor and acceptor^46^ and is taken to be 3.5 nm for Trp59-heme pair.

## Results

### Simulated and experimental time-resolved SAXS profiles are in excellent agreement

Here, we use atomically detailed simulations to probe the changes in the shape during the folding of Cyt c, a protein whose self-assembly has been extensively studied experimentally^17–19,47–49^ as well as by simulations using *C*_*α*_-based Go-models.^15,50,51^ In a previous study,^34^ the Stochastic Difference Equation in Length (SDEL), which permits generation of approximate long time folding trajectories, was used to characterize the folding energy landscape explored by Cyt Here, the folding trajectories are used to calculate the complete time-dependent scattering profiles in order to validate the simulation results. We calculated time-resolved Kratky plots for Cyt c folding by an approximate mapping of the path length to real-time (see the Methods for details and inset in Fig.1a). Figure 1a shows a comparison of the simulated *q*^2^*I*(*q*) curves at various times. Remarkably, our results are in quantitative agreement with experiments starting from the earliest time point (*t* = 160*µs*), which allows us to go further to quantify the changes during the folding of Cyt c. Representative structures from the time resolved SAXS in Figure 1a are shown in Figure 1b.

### Comparison to Hydrogen/Deuterium Exchange experiments

We also used time resolved Hydrogen/Deuterium (H/D) exchange labeling experiments^43^ as an independent data set to further validate the accuracy of our simulations. We related the changes in the H/D exchange rates to the changes in the solvent accessible surface area (SASA) of each amide hydrogen group as a function of elapsed time *t*. The time dependent protection of each amide group is monitored by the function *P*(*t, i*) that varies between (0,0.1) (details are provided in the Methods). Results are shown as a heat map in Fig. 1c. An increase in *P*(*t, i*) for residue *i* suggests a higher amide group protection, and thus, a lower H/D exchange rate. At around *t ≈* 100*µs*, the C-terminal segment 92-98 shows marked protection. N-helix protection starts at a later stage (in milliseconds). It is gratifying to note that the order of events, and timescales are in excellent agreement with ultra-fast hydrogen exchange measurements,^43^ which not only validates the simulation results but also allows us to determine the folding pathways of Cyt c using simulations. Interestingly, on the timescale, of *t ≈* 100*µs*, out of 25 residues that show protection, 17 are hydrophobic residues. The ratio 17*/*25 *≈* 0.7 is about two times the average ratio of observing a hydrophobic residue (40*/*104 *≈* 0.38) in random in Cyt c, which indicates a strong correlation between initial collapse and hydrophobicity.

### Assembly of Cyt c

In order to reveal the atomic details of the mechanism of Cyt c folding and to interpret the changes in the protection factors measured in H/D experiments, we calculated the helix content for the three major helices in the native fold of Cyt c. The time evolution of the helix content, averaged over different trajectories, are shown in Figure 2. Despite the dramatic protection in the C-terminal segment (92-98) at *t ≈* 100*µs*, there is hardly any helix formation on this timescale when canonical helix definitions are used. Helix formation occurs around 0.5 ms, about 400 microseconds after the marked increase in the amide group protections (Fig. 1c). Our simulations provide evidence for specific hydrophobic collapse concomitant with helix formation. Time dependent changes in the growth of helical content show a sequential order in the acquisition of the folded state, as found in experiments and simulations using coarse-grained models.^50,51^ Some variations in the time scale and in the order of partially folded units is evident among the different folding pathways (Figure 2).

**Figure 2:**
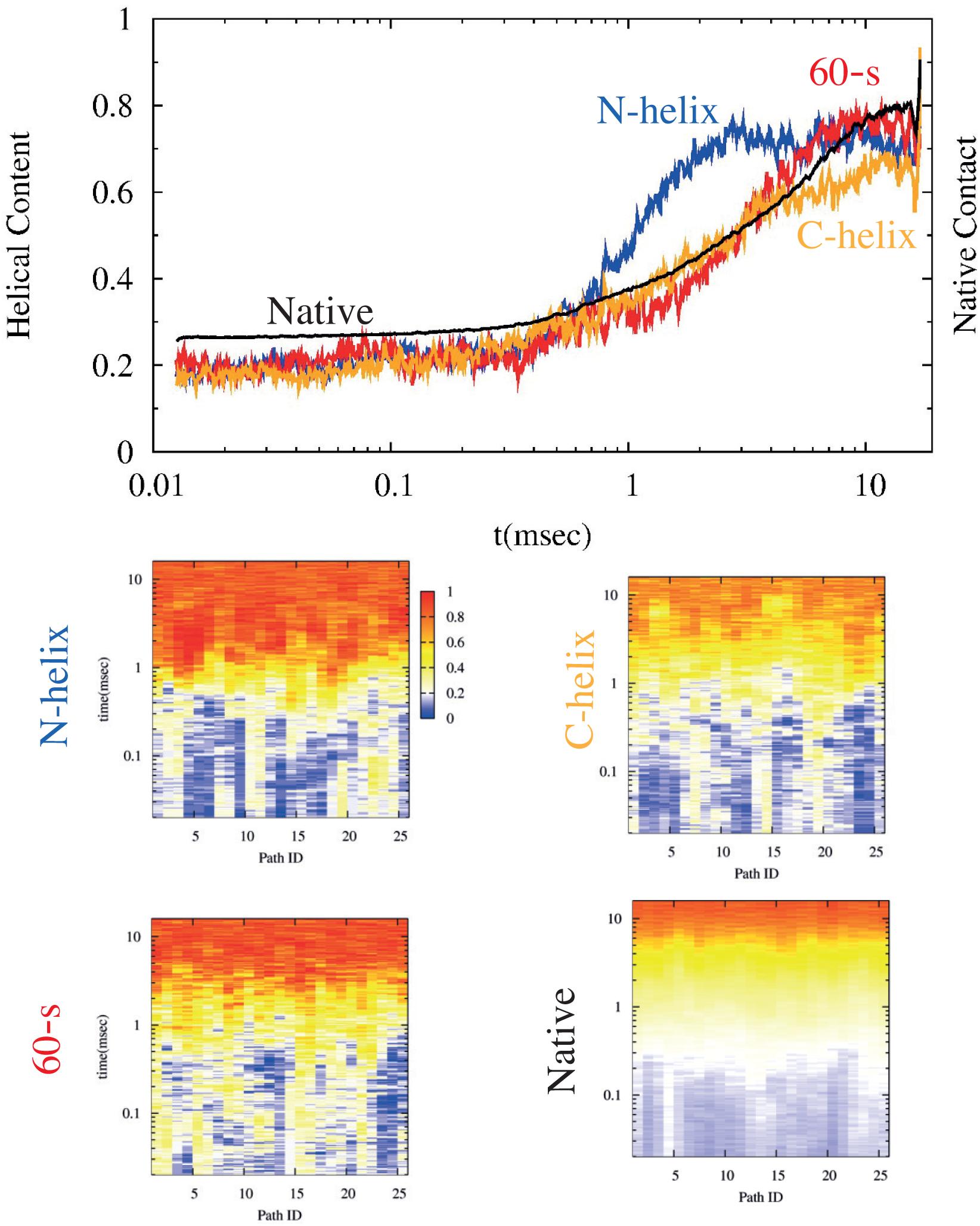
(a) Average helix content of the major helices during folding: N-helix (blue), 60’s-helix (red) and C-helix (yellow) is shown as a function of time. See Figure 1b for helix definitions. Data is averaged over all paths at a given time. Black solid line reports the native contact formation as a function of time. Two atoms of separate side chains *i* and *j* are assumed to form a contact if the distance *d*(*i, j*) *<* 5*Å*. (b) Time evolution of helix content and native contact formation of each individual SDEL trajectory. See Fig 1b for the definition of the different helical segments.

**Figure 3:**
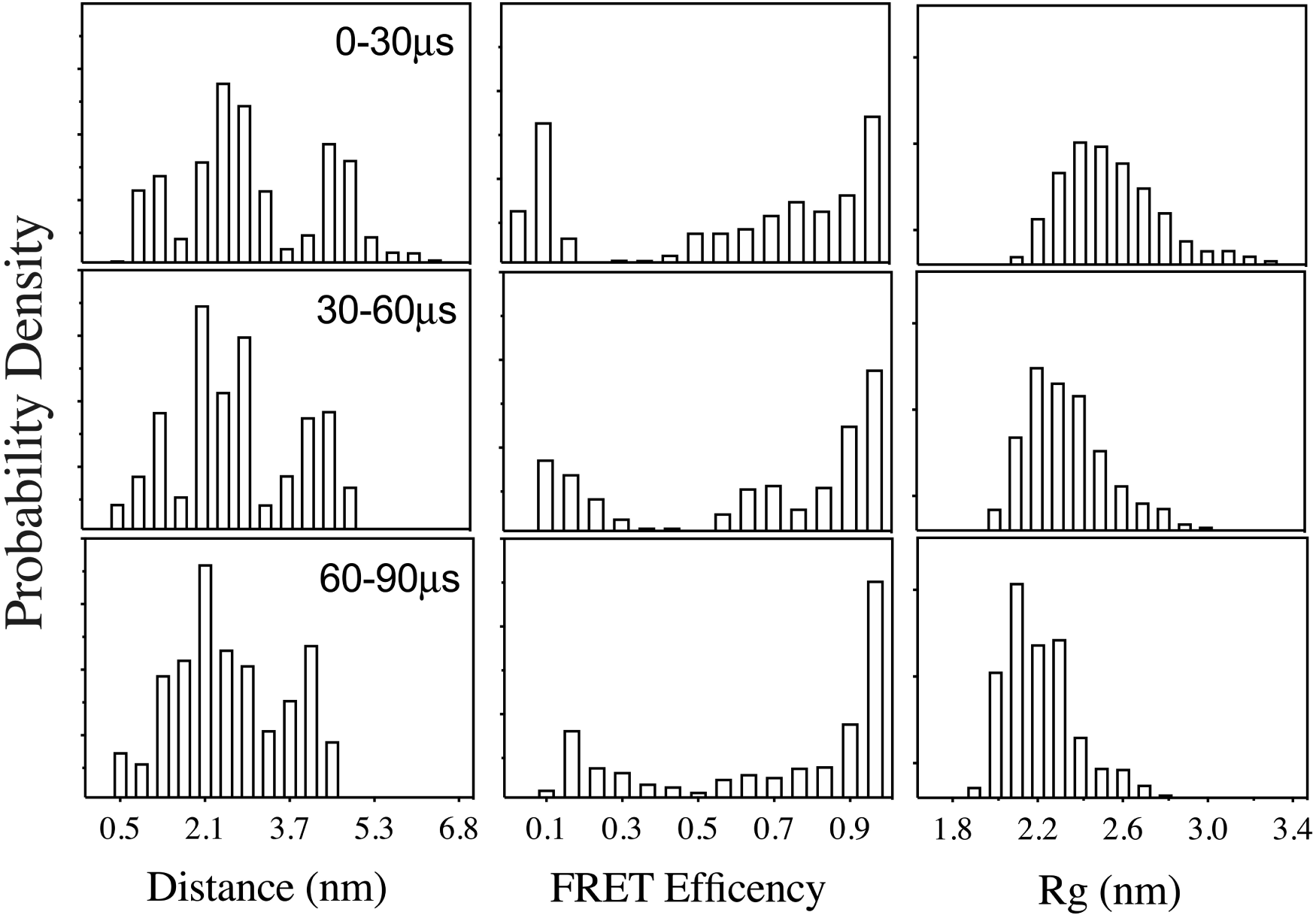
(a) Conformational change monitored for the first 90 microseconds of the Cyt C. folding. Probability density of Trp-59 Heme distance, FRET efficiency measured from the Trp59-Heme pair and radius of gyration of the whole protein are computed from folding ensemble.

**Figure 4:**
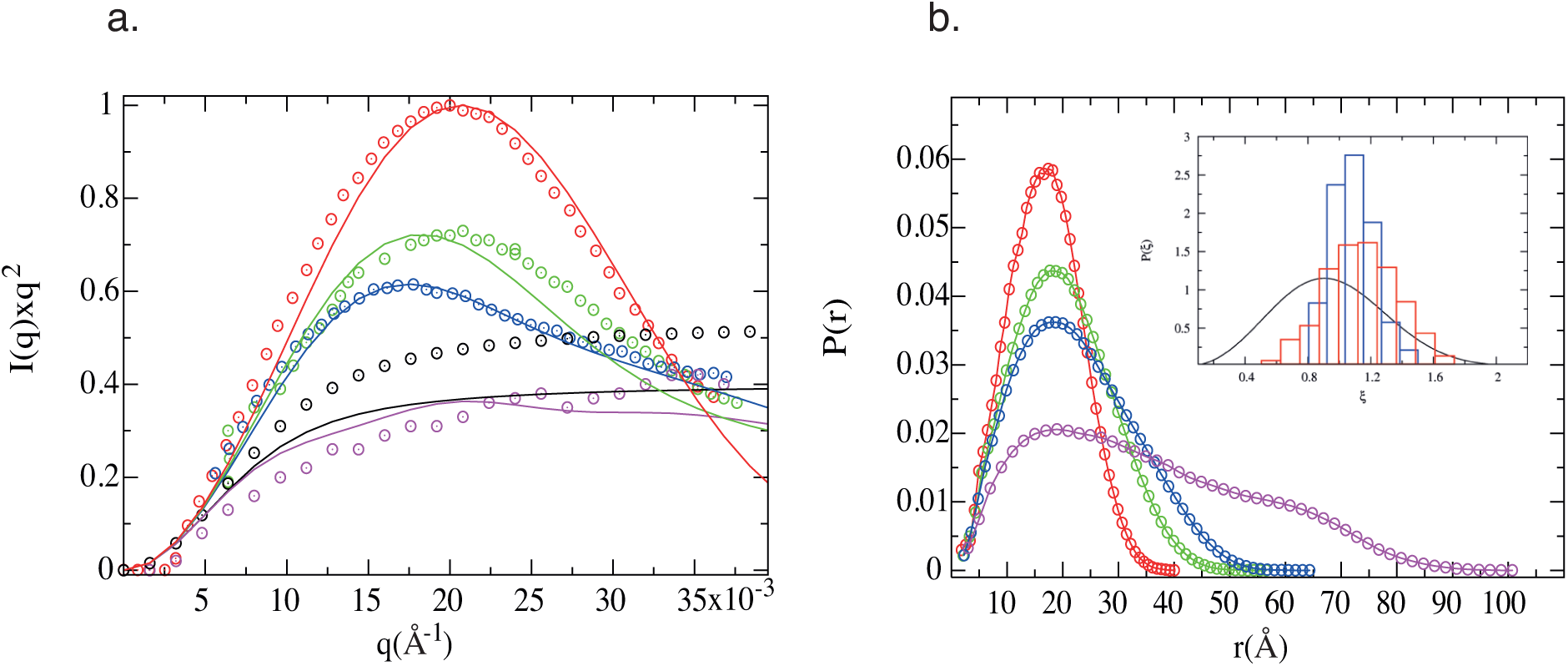
(a) Kratky plots representing the ensemble of conformations of Cyt c in different states; Native N (red), Intermediate II (green), Intermediate I (blue) and unfolded state U (purple). Circles are experimental result extracted from.^16^ Solid lines are the same quantity computed from simulations. Black circles is the Debye function when unfolding state is a random coil with *R*_*g*_ = 24.3Å, black solid line is the same function when *R*_*g*_ = 27.8Å, which is obtained from the simulation curves in Fig 1a. Based on the agreement with the Debye function using the calculated *R*_*g*_ value we surmise that *< R*_*g*_ *>* of the unfolded state is = 27.8Å. (b) Distance distribution function P(r), obtained by the inverse Fourier transform of the scattering function, computed from simulations. Color coding is the same as in Figure 1a. **Inset**. Probability distribution of 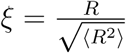 (*R* is either *R*_*g*_ or the end-to-end distance, *R*_*ee*_) for the unfolded state. Red histograms is the distribution for 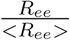, blue gives the distribution of 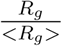 where *< R*_*g*_ *>* is the average radius of gyration. Solid line is the universal probability distribution function for variable the dimensionless v 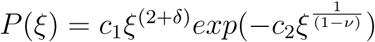 where *c*_1_ = 3.7, *c*_2_ = 1.2, 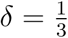, and *ν* = 0.6 for Flory random coil.

Helix formation starts with the formation of N helix, which is followed by the 60s helix formation, and is completed when the C-helix folds. These results further support the sequential mechanism for Cyt c folding, in accord with experiments^47^ and previous studies.^34,50^ From 0.5 ms to 15 ms the overall helical content increases by about 35%, which is consistent with CD experiments (see Figure 5 in^16^). We also monitored the native contact formation during folding, which shows a similar result. However, their formation is a more uniform change across the folding pathways, and occurs at a slower rate of change, suggesting a hierarchical folding after helix formation. We surmise that folding starts with hydrophobic collapse followed by secondary structure formations. Tertiary fold occurs concomitant of the secondary structure formation.

**Figure 5:**
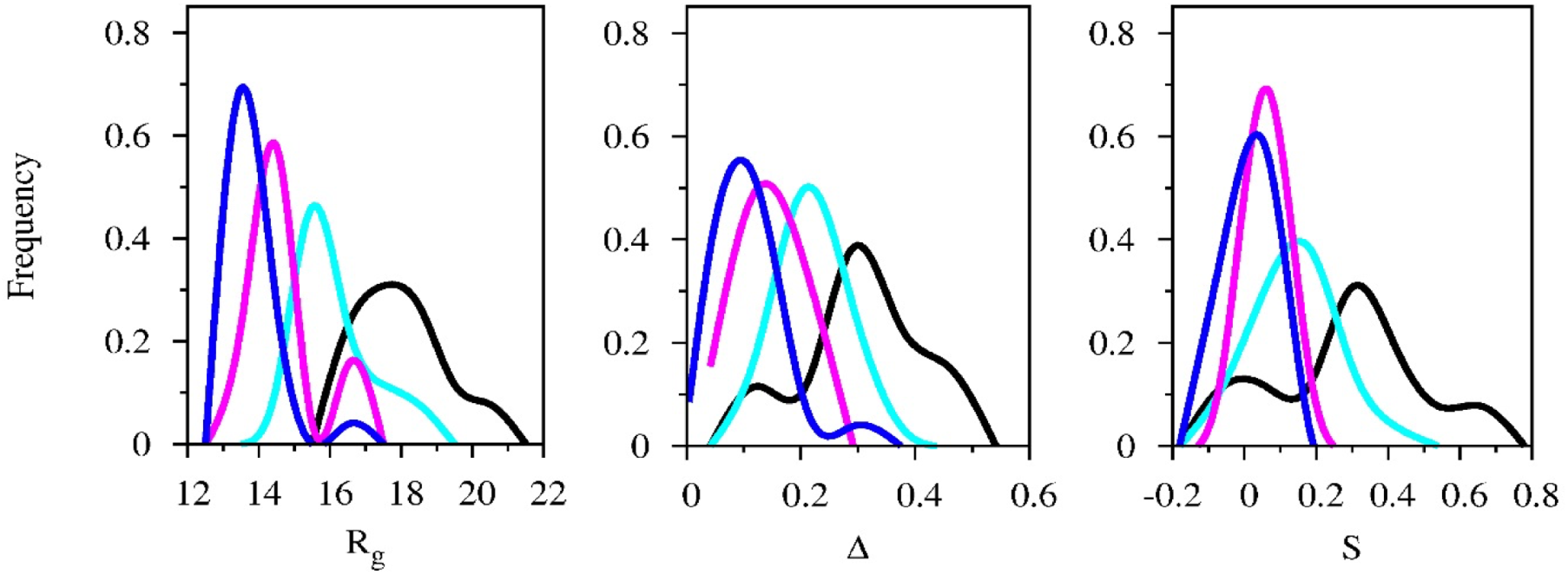
Global shape change during Cyt c folding monitored by (a) Radius of gyration (*R*_*g*_), (b) deviation from spherical symmetry (Δ) (c) structural measure of the global shape (*S*), (see SI for details). The various colors represent different stages of folding. Color coding is the same as in Figure 1a; black (*t* = 160*µs*), cyan (*t* = 550*µs*), purple (*t* = 3*ms*), blue (*t* = 6*ms*).

Despite the excellent agreement between the HX measurements and simulations at the early stages of folding,^43^ the order of folding at long times is in apparent disagreement with the interpretation of pulse labeling HX MS experiments.^52,53^ The order of events inferred from these studies indicate that the N- and C-terminal helices stably fold together and before the 60’s helix. Our amide group protection analysis also shows a strong correlation with N- and C-terminal protection while 60’s helix protection starts earlier. The discrepancy might be due to the differences in the refolding conditions, which are reflected in the refolding time scales between the two approaches: our study reports the folding time about 15ms while the experimental measurements^53^ were on *∼* 5 s. In addition, we cannot rule out the possibility that the force fields are not very accurate, especially when considering folding at long times.

### Kinetic barriers to collapse in the early stages of folding pathways

Our simulations shed light on the perennially interesting question of collapse of proteins in general, and Cyt c in particular. There are apparently conflicting views on the problem of protein collapse based on interpretation of two experiments (see^54^ for a comprehensive review). One of them measures Trp-59-heme distance from fluorescence experiments. Changes in the Trp-59 distance from the heme was used to monitor the early stages of refolding on a time scale of *≈* 100 microseconds. The bi-model nature of FRET signal reported has been correctly interpreted as arising from a kinetic barrier to the collapsed state.^55,56^ HX labeling, single molecule, and NMR measurements on the other hand, report a negligible barrier in the early stages of compaction.^57,58^ In addition, HX pulse labeling method^53^ on the slow dynamical changes reports a barrier-less transition during refolding. To reconcile these seemingly differing interpretations, we calculated the Trp59-Heme distance distributions during the early stages of folding. We also calculated the experimentally measured FRET efficiency as well as the *R*_*g*_ distribution at the early stages of folding.

The results, summarized in Figure 3 allows us to draw the following important and often under-appreciated conclusions. (i) Interestingly, the nature of the distributions depends on the probe. For example, *R*_*g*_ distributions seem to change continuously with time, suggestive of a small barrier.^59^ On the other hand, the FRET efficiency distributions show that there are two subpopulations, indicative of a barrier to collapse.^56^ The probe dependent differences in the results is a consequence of the finite size of single domain proteins, with Cyt c being no exception. (ii) It should be noted that the change in the mean value of *R*_*g*_, with respect to the unfolded states is less than about 20%, which is a common observation in a number of proteins.^60^ Our results show that it is difficult to unambiguously assess barriers to collapse in folding. In light of extensive theoretical studies, summarized elsewhere,^60^ we surmise that the barrier to collapse in Cyt c should exist^56^ but multiple probes might be needed to estimate the magnitude.

### Changes in the size of Cyt c along the folding trajectories

To elaborate on the shape changes, and characterize the missing structural details of the major states reported in,^16^ we first compare our scattering intensities with the equilibrium ensemble obtained in SAXS experiments. The structure factors for the unfolded, (U), the two intermediates (I and II), and the native (N) states are shown in Figure 4a. The distance distribution function, *P*(*r*), which is the inverse Fourier transform of *I*(*q*), for the four major folded states are shown in Figure 4b. The results of our simulations are in very good agreement with SAXS measurements for the N, I, and II states as can be seen by comparing Fig. 4b and Fig. 4 in.^16^ The simulation results reaffirmed that the largest sampled *r* value, *D*_*max*_, and the peak positions for these states. The calculated *D*_*max*_ values 40.2 Å, 59.4 Å, and 65.8 Å compare well with experimental values 39 Å, 58 Å, and 66 Å, for the states N, II and I respectively. The *R*_*g*_ values calculated using 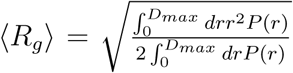 are 12.9 Å(N), 15.5 Å(II), and 17.2 Å(I) respectively. The corresponding results extracted from an initially acid denatured unfolded states in^16^ are 13.9 Å, 17.7 Å, and 21.5 Å. Taken together, we conclude that the global shape profiles obtained in experiments can be fully accounted for in our simulations.

Fig. 4 demonstrate that Cyt c collapse occurs in three major steps, as suggested previously based on lattice model simulations.^26^ In the first step, there is a substantial reduction in *R*_*g*_, reaching a value of *∼* 21 Å from about *∼* 28 Å in less then 100 *µs*. The transition from I, with *R*_*g*_ *≈* 17 Å to N occurs in two stages in our simulations with time constants of 1 and 5 ms, respectively. The two stages represent roughly I *→* II and II *→* N transitions. The absence of a direct path from U *→* N implies that the N state structure is acquired in a sequential manner, U *→* I *→* II *→* N supporting the interpretation based on the SAXS data that for *t >* 160*µs* chain collapse occurs in two major stages.

Unlike the compact states (N, II, and I), the description of the ensemble of unfolded structures (U state) show minor differences between simulation and experiment. The Kratky plot of the unfolded state is shown in Figure 4a (purple points are the experiment and purple lines are the simulation). Unfolded ensemble computed from simulation gives good agreement with the SAXS profile with an average *R*_*g*_ *≈* 28 Å for U state. The unfolded conformations sample distances nearly 2*R*_*g*_ larger than the size of the *N* state (13.9 Å). The distribution functions show that the *D*_*max*_ value for the U state extends up to *≈* 100 Å. Because *P*(*r*) for the unfolded state was not reported by Akiyama et. al,^16^ a quantitative assessment of our predictions cannot be made. However, *R*_*g*_ *≈* 28 Å from the simulation differs from the estimate of experimental, *R*_*g*_ *≈* 24.3 Å, which only about 13% smaller than the value obtained in simulations. We believe that the calculated *R*_*g*_ value is more reliable because when it is substituted into the Debye function to compute the theoretical *I*(*q*) for a random coil model there is good agreement with experiments (see the black line in Figure 4a). The experimental estimate, on the other hand, results in an unphysical extrapolation of the structure factor when the Debye function is used (black circles in Figure 4a). In addition to comparing the simulations to the Akiyama et. al^16^ experiments, we also compared them with other SAXS studies.^15,56,66^ The reports of *R*_*g*_ *≈* 30 *−* 32 Å in the unfolded state agrees well with our estimate of 28 Å. It is likely that the method of preparation of the initial unfolded state impacts the unfolded structure. In the SAXS experiments, folding was initiated by a jump in pH from 2.0 to 4.5, and *R*_*g*_ of the unfolded state was extrapolated from the time-dependent decay of *R*_*g*_ to *t* = 0. In our simulations, *R*_*g*_ is from the ensemble of conformations generated at a high temperature, which could explain the difference between the Ref.^16^ and our estimate.

One cannot rule out the possibility that the disparity between simulated and experimentally estimated value of *R*_*g*_ for the U state could also be due to the deficiencies in the use of current forcefields to describe the unfolded states. We believe that this is unlikely in this case, because the usual experience is that the all atom forcefields overestimate helicity even when the protein is unfolded. From this observation we would expect that the simulated unfolded state would be more compact when compared to experiments.^61^ Not only does the comparison show the opposite trend, the ensemble is essentially devoid of structure at *t ≈* 160*µs* (Fig. 2).

### The unfolded state is not a random coil

It is difficult to describe unambiguously whether the unfolded state can be described as a random coil using experiments alone. The mean *< R*_*g*_ *>* for the 104 residue Cyt c is predicted to be 2*N*^*ν*^ = 32 Å (*N* = 104, and the Flory exponent is *≈* 0.6). The predicted random coil value differs considerably from the calculated and SAXS measurements.^16^ In accord with this prediction, we find that the probability distribution of 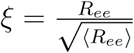 for the unfolded state, *P*(*ξ*) does not agree with the predicted universal function for random coils. This is a reflection of the potential residual structure in the unfolded state.

### Shape changes

The compaction of Cyt c, reflected in the decrease in *R*_*g*_, is also accompanied by global shape changes during the folding process (Figure 5a-c and Figure 6), which we quantify using the asphericity (Δ) and shape (*S*) parameters (see Methods for definitions). The *S* parameter is used to distinguish between prolate and oblate ellipsoids. Cyt c, at *t ≈* 160*µs*, adopts a highly elongated shape, with Δ = 0.3, and *S* = 0.3. The shape undergoes a transition to a prolate ellipsoid at *t ≈* 550*µs* with Δ = 0.2 and *S* = 0.1. A further gradual change to spherical-like shape with Δ = 0.1, and *S* = 0 occurs at 5*ms*. Figure 5 and Figure 2a show that the collapse of Cyt c occurs prior to the increase in the helical content. On the time scale of *≈* 100 *µs*, there is a substantial reduction in *R*_*g*_ from 28 Å to 20 Å (Fig. 6). At *t ≈* 0.5 ms, *R*_*g*_ decreased substantially with practically no change in the helix content. On this time scale, Cyt c is structure-less. The biphasic decrease in *R*_*g*_ also observed in recent tr-SAXS studies^56^ with time constants of *t*_1_ *≈* 45 *µs* and *t*_2_ *≈* 0.65 *ms*. These time-scales are consistent with our simulations. Upon formation of I there is an increase in the helix content, which grows rapidly when II forms in a few milliseconds, with the simultaneous formation of a well-defined core. Further consolidation occurs on the folding time scale *τ*_*F*_ *≈* 10 *ms*.

**Figure 6:**
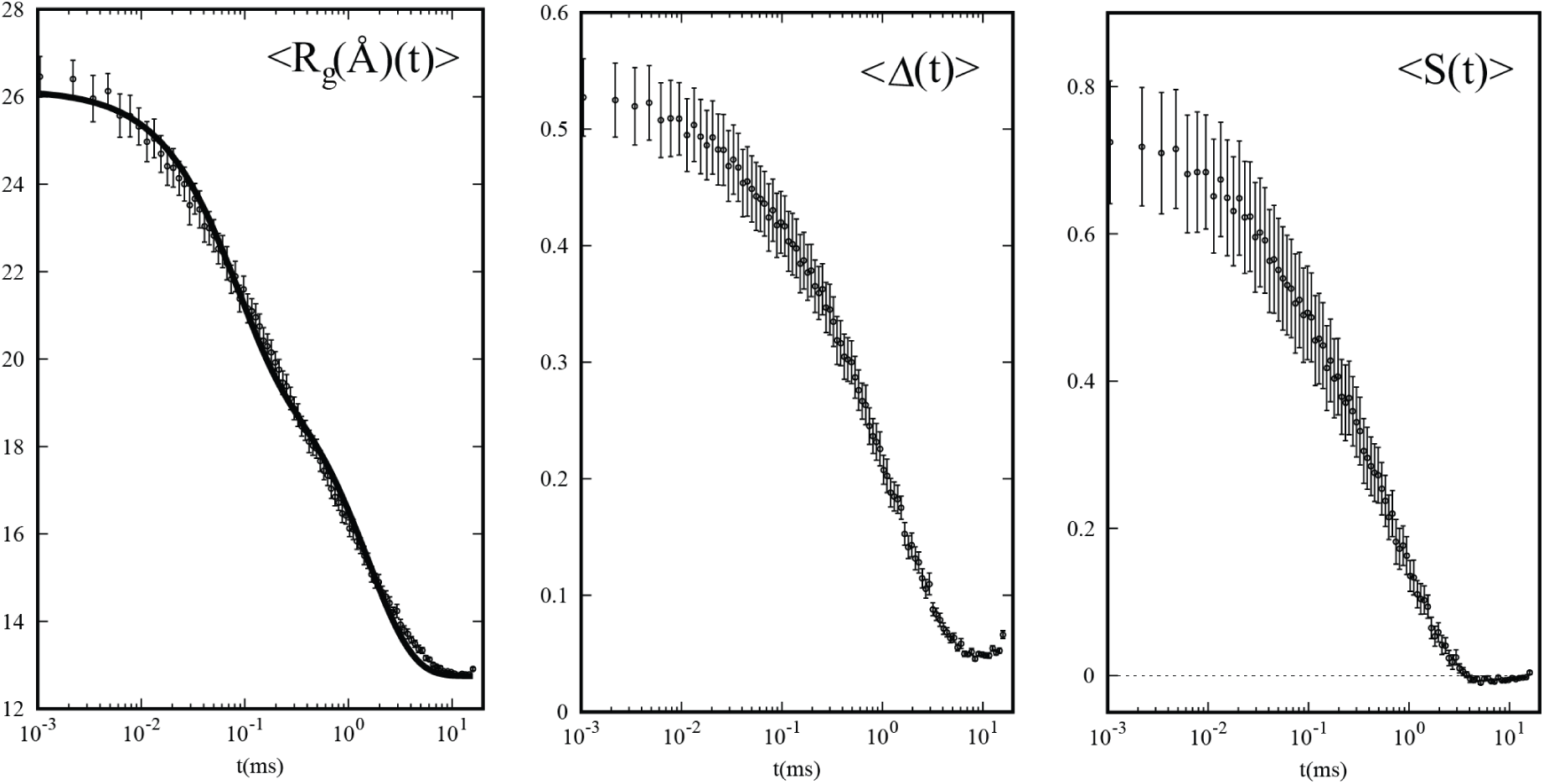
Time evolution of global shape changes monitored as average radius of gyration, asymmetry parameter, Δ and shape parameter, *S*. Circles are averages over 26 trajectories. Linear relationship between the path index and *t* is assumed to hold between 0 *< t <* 160*µs*. Decay of *R*_*g*_(*t*) is best represented by a sum of two exponential functions, 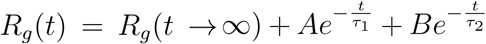 where *R*_*g*_(*t → ∞*) = 12.87Å, *A* = 6.19Å, *B* = 7.22Å, *τ*_1_ = 0.08 ms and *τ*_2_ = 1.55 ms.

## Conclusions

In this study, we used the atomically detailed folding trajectories connecting the unfolded and folded states of Cyt c to provide quantitative comparison between simulations and time-resolved SAXS experiments. The excellent agreement between the two approaches enables us to establish the structures of the key intermediates during the sequential assembly of this protein.

A few comments about the folding process of Cyt c are worth making based on our simulations: (i) The picture that emerges on how Cyt c folds is similar to that found in time resolved experiments. By simultaneously measuring the global helix content of Cyt c and the time-dependent changes in helix compaction, it was found^16^ that only upon the formation of II there is growth in the secondary structure. This interpretation is consistent with our simulations. (ii) There is a sequential order in helix organization of Cyt c, which is experimentally determined by their individual stability as studied by hydrogen exchange experiments.^47^ The global sequential folding mechanism, after collapse of Cyt c, is also in agreement with predictions based on simulations using the *C*_*α*_ based Go-models.^50^ (iii) There are a substantial number of non-native interactions that are formed in the early stages of Cyt c collapse,^34^ which are annealed subsequently, a feature first established in precisely solvable lattice models.^26^ In Cyt c, it is clear that on time scales less than about one milliseconds the structures are more native-like, which implies that specific native interactions stabilize the compact states. It appears that generically non-native interactions might play a diminishing role on time scales that exceed the collapse time scale,^62,63^ which explains the remarkable success of structure-based models in describing many aspects of the folding mechanism of small and large proteins.^12,64^ (iv) The interactions between amino acid residues are often assumed to be isotropic. However, the present simulations suggest that folding of Cyt c must also involve correct orientation of the secondary structures, which implies that angular potentials also guide self-assembly of proteins. A recent study^65^ has emphasized the importance of angle landscapes, specifically in the context of folding of cytochromes.

The dramatic changes in the shape of the protein as it folds, which occur continuously, show that by at most *t ≈* 0.55 *ms* the compaction is largely dominated by specific interactions, as opposed a global hydrophobic collapse. The residue level time-dependent accessible surface area measurements, which is a surrogate for H/D exchange protection factor, show that on *t ≈* 100 *µs*, the C-terminal helices are formed while there is disorder in the N-terminal residues. It will be interesting to perform similar simulations using our methodology in order to probe the folding process of other small globular proteins not containing the heme group.

## Acknowledgements

This work was supported by the National Foundation (CHE 16-36424) and the Collie-Welch Foundation (F-0019), NIH grant GM059796, Welch grant F-1896, and New York University Abu Dhabi faculty research grant, AD181.

## Table of Content Graphics

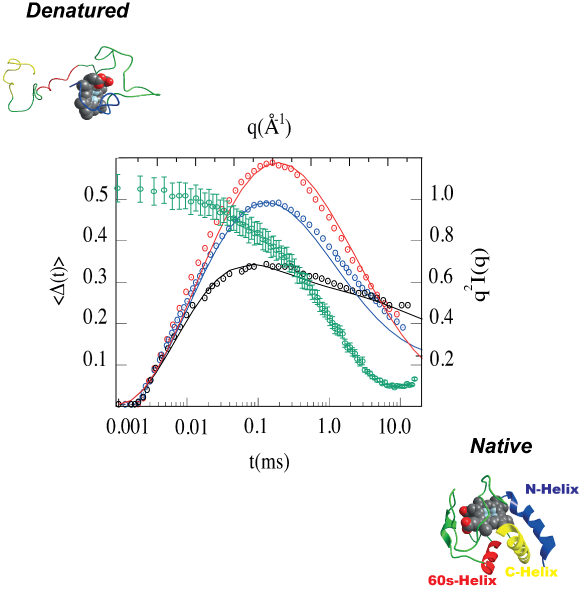

## References

(1) Schuler, B.; Eaton, W. A. Protein folding studied by single-molecule FRET. Curr. Opin. Struct. Biol., 2008, 18, 16–26.

(2) Woodside, M. T.; Garcia, C.; Blocks, M. Folding and unfolding single RNA molecules under tension. Curr. Opin. Chem. Biol., 2008, 12, 640–646.

(3) Eaton, W.A.; Munoz, V.; Hagen, S.J.; Jas, G.S.; Lapidus, L.J.; Henry, E.R.; Hofrichter, J. Fast kinetics and mechanisms in protein folding. Ann. Rev. Biophys. Biomol. Struct., 2000, 29, 327–359.

(4) Chung, H.S.; McHale, K.J.; Louis, M.; Eaton, W.A. Single-Molecule Fluorescence Experiments Determine Protein Folding Transition Path Times. Science, 2012, 335(6071), 981–984.

(5) Takahashi, S.; Kamagata, K.; Oikawa, H. Where the complex things are: single molecule and ensemble spectroscopic investigations of protein folding dynamics, Curr. Opin. Struct.Biol., 2016, 36, 1–9.

(6) Tuukkanen, A. T.; Spilotros, A.; Svergun, D. I. Progress in small-angle scattering from biological solutions at high-brilliance synchrotrons, IUCRJ, 2017, 4(5), 518–528.

(7) Shakhnovich. E., Protein folding thermodynamics and dynamics: Where physics, chemistry, and biology meet. Chem. Rev., 2006, 106, 1559–1588.

(8) Onuchic, J. N.; Wolynes, P. G. Theory of protein folding. Curr. Opin. Struct. Biol., 2004, 14, 70–75.

(9) Dill, K. A.; Ozkan, S. B.; Shell, M. S.; Weikl, T. R. The protein folding problem. Annu. Rev. Biophys., 2008, 37 289–316.

(10) Dill, K. A.; and Justin L. M. The Protein-Folding Problem, 50 Years On. Science, 2012, 338, 1042–1046.

(11) Thirumalai, D.; O’Brien, E. P.; Morrison, G.; Hyeon, C. Theoretical perspectives on protein folding. Annu. Rev. Biophys., 2010, 39, 159–183.

(12) Thirumalai, D.; Liu, Z.; O’Brien, E. P.; Reddy, G. Protein folding: from theory to practice Curr. Opin. Struct. Biol., 2013, 23, 22–29.

(13) Borgia, A.; Williams, P. M.; Clarke, J. Single-molecule studies of protein folding. Annu. Rev. Biochem., 2008, 77, 101–125.

(14) Stigler, J.; Ziegler, F.; Gieseke, A.; Gebhardt, J. C. M.; Rief. M. The complex folding network of single calmodulin molecules. Science, 2011, 334(6055), 512–516.

(15) Pollack, L.; Tate, M. W.; Darnton, N. C.; Knight, J. B.; Gruner, S. M.; Eaton, W. A.; Austin, R. H. Compactness of the denatured state of a fast-folding protein measured by submillisecond small-angle x-ray scattering. Proc. Natl. Acad. Sci., 1999, 96(18), 10115–10117.

(16) Akiyama, S.; Takahashi, S.; Kimura, T.; Ishimori, K.; Morishima, I.; Nishikawa, Y.; Fujisawa, T. Conformational landscape of cytochrome c folding studied by microsecond-resolved small-angle x-ray scattering. Proc. Natl. Acad. Sci., 2002, 99, 1329–1334.

(17) Winkler, J. R. Cytochrome c folding dynamics. Curr. Open. Struct. Biol., 2004, 8(2), 169–174.

(18) Pletneva, E. V.; Gray, H. B.; Winkler, J. R. Many faces of the unfolded state: Conformational heterogeneity in denatured yeast cytochrome c, J. Mol. Biol., 2005, 345(4), 855–867.

(19) Weinkam, P.; Pletneva, E. V.; Gray, H. B.; Winkler, J. R.; Wolynes, P. G. Electrostatic effects on funneled landscapes and structural diversity in denatured protein ensembles. Proc. Natl. Acad. Sci., 2009, 106, 1796–1801.

(20) Shaw, D. E.; Maragakis, P.; Lindorff-Larsen, K.; Piana, S.; Dror, R. O.; Eastwood, M. P.; Bank, J. A.; Jumper, J. M.; Salmon, J. K.; Shan, Y.; et al. Atomic-Level Characterization of the Structural Dynamics of Proteins. Science, 2010, 330(6002), 341–346.

(21) Vincent, A. V.; Bowman, G. R.; Beauchamp, K.; Pande, V. S. Molecular Simulation of ab Initio Protein Folding for a Millisecond Folder NTL9(1-39). J. Amer. Chem. Soc., 2010 132(5), 1526–1528.

(22) Liu, Y.; Struempfer, J.; Freddolino, P. L.; Gruebele, M.; Schulten, K. Structural Characterization of lambda-Repressor Folding from All-Atom Molecular Dynamics Simulations. J. Phys. Chem. Lett., 2012, 3(9), 1117–1123.

(23) Elber, R. Long-timescale simulation methods. Curr. Opin. Struct. Biol., 2005, 15, 151–156.

(24) Kohn, J. E.; Millett, I. S.; Jacob, J.; Azgrovic, B.; Dillon, T. M.; Cingel, N.; Dothager, R. S.; Seifert, S.; Thiyagarajan, P.; Sosnick, T. R.; Hasan, M. Z.; Pande, V. S.; Ruczinski, I.; Doniach, S.; Plaxco, K. W. Random-coil behavior and the dimensions of chemically unfolded proteins. Proc. Natl. Acad. Sci. USA, 2004, 101, 12491–12496.

(25) Dima R. I.; Thirumalai, D. Asymmetry in the shapes of folded and denatured states of proteins. J. Phys. Chem. B, 2004, 108, 6564–6570.

(26) Camacho C. J.; Thirumalai, D. Kinetics and Thermodynamics of Folding in Model Proteins. Proc. Natl. Acad. Sci. USA, 1993, 90, 6369–6372.

(27) Samanta, H. S.; Pavel, Z.; Hinczewski, M.; Hori, N.; Chakrabarti, S.; Thirumalai, D. Protein collapse is encoded in the folded state architecture. Soft Matter, 2017, 13(19), 3622–3638.

(28) Haran, G. How, when and why proteins collapse: the relation to folding. Curr. Opin. Struct. Biol., 2012, 22(1), 14–20.

(29) Ziv, G.; Thirumalai, D.; Haran, G. Collapse transition in proteins. Phys. Chem. Chem. Phys., 2009(1), 83, 83–93.

(30) Borgia, A.; Zheng, W.; Buholzer, K.; Borgia, M.; Schüler, A.; Hofmann, H.; Soranno, A.; Nettels, D.; Gast, K.; Grishaev, A.; Best, R.; Schuler, B. Consistent View of Polypeptide Chain Expansion in Chemical Denaturants from Multiple Experimental Methods, J. Amer. Chem. Soc., 2016, 138(36), 11714–11726.

(31) Hofmann, H.; Soranno, A.; Borgia, A.; Gast, K.; Nettels, K.; and Schuler, B. Polymer scaling laws of unfolded and intrinsically disordered proteins quantified with single-molecule spectroscopy. Proc. Natl. Acad. Sci., 2012, 109, 16155–16160.

(32) Ruymgaart, A. P.; Cardenas, A.E.; Elber, R. Moil-opt: Energy-conserving molecular dynamics on a gpu/cpu system. J. Chem. Theory and Computations, pages 2011, 7(10), 3072–3082.

(33) Onuriyev, A.; Bashford, D.; Case, D. A. Exploring protein native states and large-scale conformational changes with a modified generalized born model. Proteins-Structure Function and Bioinformatics, 2004, 55(2), 383–394.

(34) Cárdenas A.E.; Elber, R. Kinetics of cytochrome C folding: Atomically detailed simulations. Proteins: Structure, Function, and Genetics, 2003, 51, 245–257.

(35) Ghosh. A.; Elber, R; Scheraga, H. An atomically detailed study of the folding pathways of Protein A with the Stochastic Difference Equation. Proc. Natl. Acad. Sci., bf 2002, 99, 10394-10398.

(36) Svergun D.I.; Barberato C.; Koch M.H.J. Crysol - a program to evaluate x-ray solution scattering of biological macromolecules from atomic coordinates. J. Appl. Cryst., 1995, 28, 768–773.

(37) Resing, K. A.; Hoofnagle, A. N.; Ahn, N. G. Modeling deuterium exchange behavior of ERK2 using pepsin mapping to probe secondary structure J. Am. Soc. Mass. Spectrom., 1999, 10, 685–702.

(38) Park, I.-H.; Venable, J. D.; Steckler, C.; Cellitti, S. E.; Lesley, S. A.; Spraggon, G.; Brock, A. Estimation of hydrogen-exchange protection factors from MD simulation based on amide hydrogen bonding analysis. J. Chem. Inf. Model., 2015, 55, 1914–1925.

(39) Craig, P. O.; Lätzer, J.; Weinkam, P.; Hoffman, R. M.; Ferreiro, D. U.; Komives, E. A.; Wolynes, P. G. Prediction of native-state hydrogen exchange from perfectly funneled energy landscapes. J. Am. Chem. Soc., 2011, 133, 17463–17472.

(40) Petruk, A. A.; Defelipe, L. A.; Rodríguez Limardo, R. G.; Bucci, H.; Marti, M. A.; Turjanski, A. G. Molecular dynamics simulations provide atomistic insight into hydrogen exchange mass spectrometry experiments. J. Chem. Theory. Comput., 2012, 9, 658–669.

(41) McAllister, R. G.; Konermann, L. Challenges in the interpretation of protein H/D exchange data: a molecular dynamics simulation perspective Biochemistry, 2015, 54, 2683–2692.

(42) Mohammadiarani, H.; Shaw, V. S.; Neubig, R. R.; Vashisth. H. Interpreting Hydrogen-Deuterium Exchange Events in Proteins Using Atomistic Simulations: Case Studies on Regulators of G-Protein Signaling Proteins J. Phys. Chem. B, 2018, 122, 9314–9323.

(43) Fazelinia, H.; Xu, M.; Cheng, H.; Roder, H. Ultrafast Hydrogen Exchange Reveals Specific Structural Events during the Initial Stages of Folding of Cytochrome c. J. Amer. Chem. Soc., 2014, 136(2) 733–739.

(44) Juffer, A. H., Eisenhaber, F., Hubbard, S. J., Walther, D. Argos, P. Comparison of Atomic Solvation Parametric Sets - Applicability and Limitations in Protein-Folding and Binding. Protein Science, 1995, 4(12), 2499–2509.

(45) Mark James Abraham, Teemu Murtola, Roland Schulz, Szilárd Páll, Jeremy C. Smith, Berk Hess, Erik Lindahl GROMACS: High performance molecular simulations through multi-level parallelism from laptops to supercomputers SoftwareX, 2015, 1(2), 19–25.

(46) Förster, T. Transfer mechanisms of electronic excitation Discuss Farady Soc., 1959 7, 27.

(47) Maity, H.; Maity, M.; Krishna, M. M. G.; Mayne, L.; Englander, S.W. Protein folding: The stepwise assembly of foldon units. Proc. Natl. Acad. Sci., 2005, 102(13), 4741–4746.

(48) Krishna, M. M. G.; Maity, H.; Rumbley, J. N.; Lin, Y.; Englander, S. W. Order of steps in the cytochrome c folding pathway: Evidence for a sequential stabilization mechanism. J. Mol. Biol., 2006, 359(5), 1410–1419.

(49) Hu, W.; Kan, Z-Y.; Mayne, L.; Englander, S. W. Cytochrome c folds through foldon-dependent native-like intermediates in an ordered pathway. Proc. Natl. Acad. Sci., 2016, 113(14), 3809–3814.

(50) Weinkam, P.; Zong, C. H.; Wolynes, P. G. A funneled energy landscape for cytochrome c directly predicts the sequential folding route inferred from hydrogen exchange experiments. Proc. Natl. Acad. Sci., 2005, 102(35), 12401–12406.

(51) Craig, P. O.; Laetzer, J.; Weinkam, P.; Hoffman, R. M. B.; Ferreiro, D. U.; Komives, E. A.; Wolynes, P. G. Prediction of Native-State Hydrogen Exchange from Perfectly Funneled Energy Landscapes. J. Am. Chem. Soc., 2011, 133(43), 17463–17472.

(52) Roder, H.; Elöve G. A.; Englander, S. W. Structural characterization of folding intermediates in cytochrome c by H-exchange labelling and proton NMR Nature, 1988, 335, 700–704.

(53) Hua, W.; Kana, Z-Y.; Maynea, L.; Englander, S. W. Cytochrome c folds through foldon-dependent native-like intermediates in an ordered pathway Proc. Nat. Acc. Sci., 2016, 113, 3809–3814.

(54) Goldbeck, R. A.; Chen, E.; Kliger, D. S. Early Events, Kinetic Intermediates and the Mechanism of Protein Folding in Cytochrome c Molecular Science, 2009, 10, 1476–1499.

(55) Shastry, M.C.R.; Roder, H. Evidence for barrier-limited protein folding kinetics on the microsecond timescale Nat. Struct. Biol., 1998, 5, 385–392.

(56) Kathuria, S. V.; Kayatekin, C.; Barrea, R.; Kondrashkina, E.; Graceffa, R.; Guo, L.;, Nobrega, R. P.; Chakravarthy, S.; Matthews, C. R.; Irving, T. C.; Bilsel. O. Microsecond Barrier-Limited Chain Collapse Observed by Time-Resolved FRET and SAXS J. Mol. Biol., 2014, 426, 1980–1994.

(57) Sosnick, T.R.; Mayne, L.; Englander, S.W. Molecular collapse: the rate-limiting step in two-state cytochrome c folding. Proteins, 1996, 24, 413–426.

(58) Qiu, L.; Zachariah, C.; Hagen, S.J. Fast chain contraction during protein folding: “Foldability” and collapse dynamics Phys. Rev. Lett., 2003, 90, 168103.

(59) Thirumalai, D. From Minimal Models to Real Proteins: Time Scales for Protein Folding Kinetics. J. Phys. I France, 1995, 5, 1457–1467.

(60) Thirumalai, D.; Samanta, H. S.; Maity, H.; Reddy G. Universal Nature of Collapsibility in the Context of Protein Folding and Evolution. Trends in Biochem. Sci., 2019, 44, 675–687.

(61) Piana, S.; Klepeis, J. L.; Shaw, D.E. Assessing the accuracy of physical models used in protein-folding simulations: quantitative evidence from long molecular dynamics simulations. Curr. Opin. Struct. Biol., 2014, 24, 98–105.

(62) Camacho C. J.; Thirumalai, D. Modeling the Role of Disulfide Bonds in Protein-Folding Entropic Barriers and Pathways Proteins: Structure, Function, and Genetics, 1995, 22(1), 27–40.

(63) Borgia, A.; Kemplen, K. R.; Borgia, M. B.; Soranno, A.; Shammas, S.; Wunderlich, B.; Nettels, D.; Best, R. B.; Clarke, J.; Schuler, B. Transient misfolding dominates multidomain protein folding, Nat. Comm., 2015, 6, 8861–7.

(64) Whitford, P.; Sanbonmatsu, K. Y.; Onuchic, J. Biomolecular dynamics: order–disorder transitions and energy landscapes Rep. Prog. Phys., 2012, 75, 076601.

(65) Kozak, J. J; Harry B. Gray, H. B.; Garza-López, R. A. Funneled angle landscapes for helical proteins J. of. Inorg. Biochem., 2020, 208, 1–13

(66) Segel, D. J.; Fink, A.L.; Hodgson, K.O.;, Doniach, S. Protein Denaturation: A Small-Angle X-ray Scattering Study of the Ensemble of Unfolded States of Cytochrome c J. Phys. Chem. B, 2018, 122, 9314–9323.

